# Revealing the impact of *Pseudomonas aeruginosa* quorum sensing molecule 2’-aminoacetophenone on human bronchial-airway epithelium and pulmonary endothelium using a human airway-on-a-chip

**DOI:** 10.1101/2025.03.21.644589

**Authors:** Shifu Aggarwal, Arijit Chakraborty, Vijay Singh, Stephen Lory, Katia Karalis, Laurence G. Rahme

## Abstract

*Pseudomonas aeruginosa* (PA) causes severe respiratory infections utilizing multiple virulence functions. Our previous findings on *PA* quorum sensing (QS)-regulated small molecule, 2’-aminoacetophenone (2-AA), secreted by the bacteria in infected tissues, revealed its effect on immune and metabolic functions favouring a long-term presence of *PA* in the host. However, studies on 2-AA’s specific effects on bronchial-airway epithelium and pulmonary endothelium remain elusive. To evaluate 2AA’s spatiotemporal changes in the human airway, considering endothelial cells as the first point of contact when the route of lung infection is hematogenic, we utilized the microfluidic airway-on-chip lined by polarized human bronchial-airway epithelium and pulmonary endothelium. Using this platform, we performed RNA-sequencing to analyse responses of 2-AA-treated primary human pulmonary microvascular endothelium (HPMEC) and adjacent primary normal human bronchial epithelial (NHBE) cells from healthy female donors and potential cross-talk between these cells. Analyses unveiled specific signaling and biosynthesis pathways to be differentially regulated by 2-AA in epithelial cells, including HIF-1 and pyrimidine signaling, glycosaminoglycan, and glycosphingolipid biosynthesis, while in endothelial cells were fatty acid metabolism, phosphatidylinositol and estrogen receptor signaling, and proinflammatory signaling pathways. Significant overlap in both cell types in response to 2-AA was found in genes implicated in immune response and cellular functions. In contrast, we found that genes related to barrier permeability, cholesterol metabolism, and oxidative phosphorylation were differentially regulated upon exposure to 2-AA in the cell types studied. Murine *in-vivo* and additional *in vitro* cell culture studies confirmed cholesterol accumulation in epithelial cells. Results also revealed specific biomarkers associated with cystic fibrosis and idiopathic pulmonary fibrosis to be modulated by 2-AA in both cell types, with the cystic fibrosis transmembrane regulator expression to be affected only in endothelial cells.

The 2-AA-mediated effects on healthy epithelial and endothelial primary cells within a microphysiological dynamic environment mimicking the human lung airway enhance our understanding of this QS signaling molecule. This study provides novel insights into their functions and potential interactions, paving the way for innovative, cell-specific therapeutic strategies to combat *PA* lung infections.

## Introduction

Pathogens have evolved diverse mechanisms to manipulate host cell functions, facilitating their survival and evasion of the host’s immune response. Bacterial quorum sensing (QS), a cell density-dependent conserved system, via the synthesis and secretion of signaling molecules, coordinates various virulence activities in Gram-negative and Gram-positive bacteria and plays an important role in the modulation of immune and metabolic functions (1–6)

*Pseudomonas aeruginosa (PA)*, a recalcitrant ESKAPE pathogen, is notorious for its rapid development of antibiotic resistance and its ability to cause acute and chronic infections. This opportunistic pathogen can infect diverse tissues, including the lungs, as a dominant respiratory pathogen. It employs multiple mechanisms to target and damage epithelial and endothelial cells, including adherence and invasion, secretion of various exotoxins, and inhibition of angiogenesis (7, 8). *PA* can adhere to and invade these cells and survive intracellularly for extended periods, leading to progressive endothelial and epithelial tissue damage (7–9). This pathogen causes acute pneumonia and chronic infections in individuals with compromised lung defenses, such as people with cystic fibrosis (CF), those with bronchiectasis, chronic obstructive pulmonary diseases, and critically ill patients on mechanical ventilation or otherwise immunocompromised patients (10–13). A commonality among these conditions is the impairment of the lung’s innate immunity, characterized by dysfunction of the mucociliary escalator, mucus accumulation, damage to the lung epithelial barrier, and persistent inflammation. These factors collectively heighten susceptibility to infections, disrupt local immune response, and hinder effective treatment (8).

*PA* QS systems, LasR/I, RhlR/I, and MvfR/PqsABCDE are responsible for the synthesis of various low molecular weight signaling molecules to regulate its virulence (14), with several shown to modulate host immune response (2, 3, 15–17). Specifically, the MvfR/PqsABCDE system, also called Pqs, regulates the synthesis of the 4-hydroxy-2-alkylquinolines (HAQs), and the non-HAQ molecule, 2’-aminoacetophenone (2-AA) (18–20). 2-AA is synthesized and secreted in *P. aeruginosa*-infected human tissues and is considered a promising breath biomarker in patients with *PA* infection (2, 3, 19, 21, 22). Small molecules secreted by bacteria at the infection site can enter the bloodstream (23, 24). Once in circulation, these molecules may rapidly reach the pulmonary capillaries, where their first point of contact is the endothelial cells lining the capillary walls (25). Endothelial cells can allow translocation and/or raise a direct response to these bacterial molecules, thereby influencing epithelial cell responses, although the cascade of events resulting in epithelial infection has not been characterized in detail. Independently of the dissemination of the bacterial-produced small molecules, infections disseminating via the hematogenous route are common in immunocompromised patients (26, 27).

Our previous studies in *PA* uncovered unique immunometabolic mechanisms by which 2-AA mediates mutual pathogen-host adaptation (2, 15, 21, 28). We identified how 2-AA induces host epigenomic reprogramming via metabolic derangement and rewires immune and metabolic functions to enable tolerance to persistent infection, while rescuing the mortality of the infected animals (2, 3, 15–17). 2-AA-induced immune tolerization in macrophages leads to metabolic perturbations in mitochondrial functions (3). It impacts the autophagic machinery and lipid biosynthesis to sustain *PA’s* presence in macrophages (3). In skeletal muscle, 2-AA also compromises mitochondrial functions (1, 29), leading to increased oxidative stress and subsequent apoptosis (28). Despite the described multifaceted role of this small molecule, its effect on pulmonary function and physiology remains poorly understood.

Microphysiological systems leveraging technologies for organ-on-chip engineering have emerged in the last few years (30–32). Studies have shown that with these microengineered devices, critical elements of human organ physiology, cell specific changes associated with disease states, and mechanisms driving therapeutic responses to clinically relevant drug exposures can be recapitulated with high fidelity (33–37). Commercially available Lung-Chip platforms provide an engineered “microtissue” emulating the cell-cell interface and allowing for controlled mechanical forces such as stretch and shear stress, flow, cell exposure to *in vivo* relevant biochemical cues, and signals (38–40) and readouts with the experimentally desired spatiotemporal resolution. It enables detailed molecular, biochemical, and metabolic studies on lung tissue cells maintained in an epithelial/endothelial interface supported by a human tissue-relevant extracellular matrix, allowing for an *in vivo* relevant barrier formation. This interface is critical in preventing the non-specific migration of cells through the membrane pores, enabling functional maturation of the cells within the chip, and closely recapitulating the vascular molecule transfer to the epithelium. Given the increasing concern over bacterial infections and their effects on pulmonary health, understanding the mechanisms underlying epithelial-endothelial cell interactions during microbial interactions with the host or the impact of secreted bacterial products is vital in developing new therapeutics and identifying disease biomarkers.

In this study using the cutting-edge human airway-on-a-chip platform, we interrogated the 2-AA impact on the human primary pulmonary microvascular endothelial cells (HPMEC) and normal human bronchial epithelial (NHBE) cells from healthy donors, considering pulmonary endothelial cells as the first point of contact. This may occur in *PA-*disseminated infections when the route of lung infection is through the bloodstream. *PA* actions on endothelial cells compromise vascular integrity and play a significant role in the pathogenesis of infections caused by this versatile pathogen, which potentially contributes to bacteremia in compromised patients (41). Our data uncovered multiple common and cell-specific 2-AA-mediated responses affecting various signaling pathways and genes, providing novel insights into the effects of this molecule that can inform the development of new therapeutics to mitigate the effects of this pathogen.

## Materials and Methods

### Activation of the airway-on-a-chip platform

The human airway-on-a-chip platform used in this study is the 2-channel microfluidic Chip-S1 Organ Chip device fabricated by Emulate Inc. It consists of two parallel channels: the mucociliary airway epithelium at the top and a microvascular endothelium at the bottom channel, separated by a human tissue-relevant extracellular matrix. The PDMS membrane was activated one day prior to introducing primary human lung microvascular endothelial cells (HPMEC) and normal human bronchial epithelial cells (NHBE) from healthy female donors using ER1 (Emulate reagent 1) and ER2 (Emulate Reagent 2) reagents (Emulate, 10465). A total of 50 μL (0.5 mg/mL) ER1 solution resuspended in ER2 was introduced in the top and bottom channels through the corresponding inlets using a 200μL micropipette. Chips were incubated under UV light for 10 minutes (min), followed by 3 min incubation at room temperature and another 10 min of UV treatment. Subsequently, both channels were washed with ER2, then with cold 1X DPBS (Dulbecco’s Phosphate Buffered Saline) and coated overnight at 37°C with 50 μL solution of 50 μg/mL Fibronectin (Corning, 354008), 50 μg/mL Laminin (Millipore Signa, 05-23-3703 1MG) and 100 μg/mL Collagen I (Advanced BioMatrix (50-360-233) added in the inlets of the top and bottom channel.

### Establishment of the HPMEC and NHBE cell cultures in the airway-on-a-chip and treatment with 2-AA

Following the overnight coating, chips were washed twice with 1X PBS. The endothelial cells (HPMEC) (Millipore Sigma, C-12281), at a concentration of 6 ×10^6^ cells/mL (∼30 μL) diluted in Endothelial Cell Growth Medium MV 2 (Promocell, C-22221 and Growth Medium MV 2 Supplement Pack C39221), were introduced through the inlets of the bottom channel. Chips were inverted, and cells were left to adhere for 2 hours at 37°C,5% CO_2_. Chips were then inverted back and epithelial (NHBE) cells (Lonza, CC-2570) at a concentration of 3x10^6^ cells/mL (approximately 40 μL), diluted in epithelial growth medium (PromoCell, C-21260, and Supplement Pack Airway Epithelial Cell GM C39160), were introduced into the inlet of the top channel.

To remove the non-adherent cells the following day, the upper channel was washed gently with the epithelial growth medium and the bottom channel with the endothelial cell growth medium MV 2. The chips were attached to Pod™ Portable Modules (Emulate Inc., 10153) and connected to the automated cell culture system Zoë™ at a flow rate of 30 μL/h in both channels to establish a monolayer for 7 days. Culture media was refreshed every two days. From day 8 through day 14, the epithelial channel was switched to Air Liquid Interface (ALI) by removing the media from the top channel to allow cell epithelial cell polarization. The bottom channel was continuously perfused with a fresh medium mimicking human vasculature using the PneumaCult™-ALI Medium (Stemcell, 05002) supplemented with 10 ng/mL VEGF, 1 µg/mL Vitamin C was added, and the flow rate was set at 40 µl/hr. At day 14, 20 μM of the 2-AA compound (Sigma, A37804) was added in the ALI medium (bottom channel) with a continuous flow for 20 hrs. After treatment, the medium was removed from the bottom channel, and chips were used to extract or stain RNA.

### RNA isolation from the chips

The top and bottom channels of the chip were lysed, NHBE and HPMEC cells were collected separately. Both channels were rinsed once with 200 μL of ice-cold PBS. Subsequently, we blocked the inlet and outlet of the channel opposite the one containing the cells of interest using empty 200 μL tips. We gently rewashed the channel of interest with 200 μL of ice-cold PBS. Once the washing step was completed, the PBS was gently aspirated from the channel, leaving it dry. The outlet port of the channel of interest was blocked with an empty 200 μL tip. The tip was not pushed entirely against the bottom of the channel to allow for a smooth flow of lysis buffer in and out of the pipette tip. Then, we introduced 70 μL of lysis buffer (Qiagen, 74004) into the channel of interest (through the inlet port) using a 200 μL tip. The cell lysate was collected in an RNase-free Eppendorf Tube® and placed on ice or stored at -80°C. RNA extraction from the cells was isolated using the RNeasy Micro Kit (Qiagen, 74004). Genomic DNA was removed using the RNase-Free DNase Set 1500 units (Qiagen, 1023460).

### RNA sequencing and data analysis

RNA isolated from untreated and 2-AA treated (n=2) HPMEC and NHBE cells were sequenced using an Illumina platform (MGH core facility). rRNA from the samples was depleted (ribosomal depletion Kit), and sequencing was performed (paired-end reads of 100 bp) using an Illumina Hiseq2500 platform.

The RNA sequencing data was analyzed using the cloud-computing server of Galaxy (usegalaxy.org), which is an open-source, web-based platform for next-generation sequencing analysis (42). The paired-end reads obtained were quality-controlled using FastQC (Galaxy version 0.74). Trimming of the 3’adaptor and the low-quality read were removed using the Cutadapt tool (Galaxy version 4.8). The trimmed paired-end reads were joined using the concatenate tool (Galaxy version 9.3). The reads were mapped to Homo sapiens (release 38) reference sequence (GRCh38) using STAR alignment (galaxy version 2.7.11a). Genome annotations for the human reference were obtained from the UCSC genome browser gateway. Reads mapped to the GRCh38 were counted using featureCounts (Galaxy version 2.0.3) with default parameters.

Differential gene expression analysis was implemented using DeSeq2 version (1.40.2) with default parameters. Genes with an increase or decrease of at least 0.5 or -0.5 log2 fold change and a maximum p-value of 0.05 were considered as significantly regulated genes. Volcano plots were generated using graphpad prism software. Heatmaps were created using Galaxy Heatmap2 (version 3.1.3.1). The Reactome pathway enrichment analyses of the significantly regulated genes were visualized by the DAVID webserver (https://david.ncifcrf.gov).

### Airway-on-a-chip staining

Following media removal from both channels, chips were stained with 1 μM of working solution of mitotracker dye (Fisher, M7512) and kept for 15 mins at 37 °C with 5% CO2, protected from light. The chips were fixed with 4% paraformaldehyde (PFA) in PBS and kept for 30 mins in the dark at room temperature with 200 μl tips in the ports. Chips were washed once with 1X PBS for 5 mins, followed by a 10 min wash with 1X PBST. The chips were stained with filipin (Sigma, SAE0087; 1:2000 dilution) and nuclear green LCS1 (Abcam, ab138904; 1:4000 dilution) in PBS for 1 hr at room temperature in the dark. Chips were rinsed three times with 1X PBS, and visualization was done by confocal microscopy (Nikon ECLIPSE Ti2; NIS-Elements 5.21; Nikon Instruments Inc., Tokyo, Japan). The fluorescence intensity of the mitotracker (red channel), Filipin (blue channel), and DAPI (green channel) was measured using the ImageJ software. The fluorescence intensity arbitrary units were calculated by normalizing the mitotracker with LCS1 and Filipin with the corresponding LCS1 stain.

### Quantification of cholesterol from the chip effluent

Effluent from the endothelial channel, which is maintained at the air-liquid interface (ALI), was collected before treatment (0 hr) and after 20 hrs of treatment with either 20 μM 2-AA or PBS. Following the manufacturer’s instructions, the cholesterol levels in the collected effluents were quantified using the Amplex Red cholesterol assay kit (Fisher, A12216). Cholesterol concentrations in the 2-AA-treated and the PBS-treated groups were normalized using the standard curve generated from known cholesterol concentrations.

### Mice treatment and lung tissue stainings

Six-week-old female C57BL/6 mice received a single intraperitoneal (IP) injection of 100 μl of 2-AA in PBS (6.75 mg/kg body weight). The control group received 100 μl of PBS. After 4 days, the mice were euthanized, and the lungs were perfused with normal saline via the trachea. Lungs were inflated with 10% neutral buffered formalin, the trachea was tied off, and the lung was removed for overnight fixation in formalin. Fixed lungs were dehydrated and embedded in paraffin wax using disposable base molds (Fisher HealthCare; #22363553). Consecutive 5 μm sections were generated from the lung tissue blocks using a Leica RN2025 microtome. Lung sections were mounted onto glass microscopic slides and stored at room temperature until use. Slides were deparaffinized and rehydrated using the 1X antigen retrieval solution. Sections were washed twice with 1X PBS (pH=7.4) and incubated with 10% goat serum for 1 hr at room temperature to block non-specific binding.

To assess cholesterol biosynthesis, we used the anti-HMGCR (ab242315, Abcam; 1:200 in 10% goat serum) primary antibody overnight at 4°C, which was protected from light. The slides were washed three times with 1X PBS and incubated with fluorescently labeled secondary antibody (Abcam, ab 175700; 1:200 in 10% goat serum) for 1 hr at room temperature. The tissue nuclei were counterstained with DAPI (Sigma, D9542; 1:10000) for 5 mins at room temperature, followed by three washes with 1X PBS (pH=7.4). Stained tissue sections were examined using a confocal microscope (Nikon ECLIPSE Ti2; Nikon Instruments Inc., Tokyo). Images were captured from at least six random fields per section. Fluorescence intensity was quantified using the Image J software.

For the hematoxylin-eosin (H&E) staining, the lung tissue sections (5 μm) were stained with H&E and, dehydrated through a series of graded alcohol and cleared in xylene. Finally, the sections were mounted with DPX (dibutyl phthalate in xylene) mounting medium. The mounted sections were visualized and imaged using an Echo Rebel microscope.

### Filipin Staining on chamber slides

NHBE and HPMEC cells were grown to 95% confluency on chamber slides (Ibidi, 80826). Cells were treated with PBS (control) or 200 μM 2-AA for 24 hrs at 37 °C, 5% CO2. After treatment, cells were washed twice with PBS (pH=4.7) and fixed with 4% paraformaldehyde for 15 mins at room temperature. A filipin working solution was prepared by diluting filipin III (Sigma, SAE0087) to 1:2000 in PBS. Cells were stained with the filipin working solution for 1 hr at room temperature in the dark. Stained cells were visualized by confocal microscopy (Nikon ECLIPSE Ti2; NIS-Elements 5.21; Nikon Instruments Inc., Tokyo, Japan). Images were captured from at least six random fields per replicate. The fluorescence intensity of the green and blue channels was measured using the ImageJ software. The fluorescence intensity arbitrary units were calculated by normalizing the blue fluorescence of Filipin with the corresponding green fluorescence values representing LCS1 stain.

### Quantification of total cholesterol in HPMEC and A549 cell lines

The HPMEC cells were cultured in T-75 flask using Endothelial Cell Growth Medium MV 2 (Promocell, C-22221 and Growth Medium MV 2 Supplement Pack C39221) with 1X penicillin/streptomycin (Gibco). Human lung epithelial A549 cells were maintained in T-75 flasks containing RPMI-1640 medium (Gibco) supplemented with 10% heat-inactivated FBS (endotoxin-free Certified FBS; Invitrogen), 1X penicillin/streptomycin (Gibco). Both cell lines were maintained at 37 °C with 5% CO2. Cells were used between passages 3 and 4.

HPMEC and A549 cells were seeded in 100 mm Petri dishes 95% confluency. Cells were treated with 200 μM of 2-AA for 24 hrs, with PBS-treated cells as control. Before and after treatment (0 hr and 24 hr), the supernatant was collected for extracellular cholesterol measurement. The cells were collected after the treatment to measure the total cell cholesterol. The cells were centrifuged at 2000 rpm for 5 mins, and the supernatant was discarded. The cell pellet was resuspended in ice-cold RIPA lysis buffer (Cell Signalling Technology, USA) containing a protease inhibitor cocktail (Millipore 1183580001) and was kept on ice for 1 hr. The cells were centrifuged at 5000 rpm for 15 minutes, and the supernatant was collected to quantify the total cell cholesterol. Extracellular cholesterol levels and total cell cholesterol were quantified using the Amplex Red cholesterol assay kit (Fisher, A12216) following the manufacturer’s instructions. Cholesterol concentrations were normalized with the standard curve created using the known cholesterol concentrations.

### Statistical analysis

All statistical analyses were performed using GraphPad Prism 9 software except for RNA-sequencing analysis. Wherever applicable, at least four or five independent experiments were performed. Data was analyzed using a two-tailed t-test and ordinary one-way analysis of variance (ANOVA) with Tukey’s post hoc test. For all experiments, p-values of <0.05 were considered significant.

## Results

### 2-AA impacts the gene expression of human pulmonary microvascular endothelium and bronchial-airway epithelium

Employing the human airway-on-chip platform, we evaluated the impact of 2-AA on human pulmonary microvascular endothelium (HPMEC) and normal human bronchial-airway epithelium (NHBE) cells. Figure 1A describes the platform, the timeline, and the steps of preparation and 2-AA treatment of these primary human cells with NHBE cells in the top channel and HPMEC cells in the bottom channel separated by a porous PDMS membrane.

**Figure 1:**
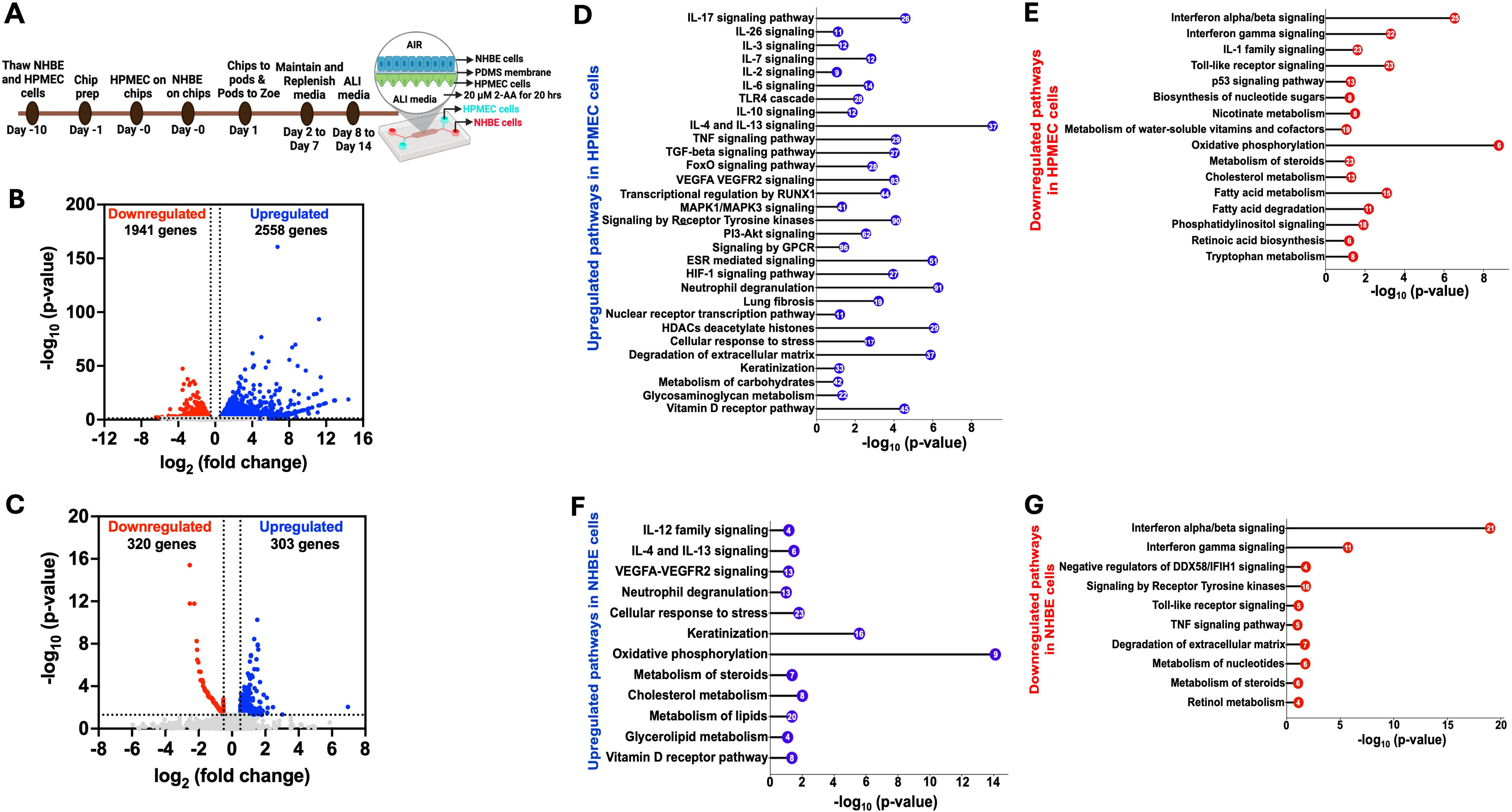
Airway-on-chip RNA-sequencing analysis of 2-AA-treated endothelial and epithelial cells reveals multiple pathways to be differentially regulated by this quorum sensing molecule. (**A**) Overview of the timeline of preparation and treatment of 2-AA of human pulmonary microvascular endothelial (HPMEC) and normal bronchial epithelial (NHBE) cells in a lung-on-chip setting. Volcano plot of the differentially expressed genes (DEGs) in HPMEC (**B**) and NHBE (**C**) cells after 20 hrs of continuous flow of 20 µM 2-AA. The blue dots represent genes that are upregulated (log_2_fold change>0.5), red dots for downregulated (log_2_fold change <-0.5), and grey dots for genes that are not significant. The two vertical dashed lines indicate log_2_fold change>0.5 and log_2_fold change <-0.5. The horizontal dashed line indicates the significance of -log_10_ (p-value) =1.3. (D-G) Horizontal lollipop plots showing Reactome pathway enrichment analysis of upregulated and downregulated pathways from the differentially regulated DEGs in human HPMEC and NHBE cells and compared to untreated cells with -log_10_(p-value) =1.3 on the x-axis. The numbers in the circles represent the count of genes for each pathway. Deseq2 determined the log2fold change values used in volcano plots. The -log_10_ (p-value) of 1.3 corresponds to p<0.05 as determined by Deseq2. The Reactome pathway enrichment analysis was done using the DAVID web server. The RNA sequencing analysis was done from two independent experiments n=2.

To address the impact of 2-AA on endothelial cells as the first point of contact when the route of lung infection is through the bloodstream, we delivered 2-AA (20 µM) to the HPMEC channel and exposed cells to continuous 2-AA flow for 20 hrs. In this setting, 2-AA can also reach the NHBE cells through the porous membrane. Continuous PBS flow in the non-2-AA-treated chips was used as a control. Following 20 h post-exposure to 2-AA or PBS, RNA was extracted from the cells of each channel individually, and RNA sequencing was performed. Principal component analysis (PCA) of the RNA-seq data revealed distinct differences between the 2-AA-treated and PBS-treated cells and between HPMEC and NHBE 2-AA-treated cells (Figures S1A and S1B). Transcriptomic profiling showed that 2-AA significantly impacts the transcription of multiple genes in both endothelial and epithelial cells with altered expression of 4499 genes in HPMEC and 623 genes in the NHBE cells. In HPMEC, 2558 genes were upregulated, and 1941 genes were downregulated (Figure 1B), while in NHBE cells, 303 genes were upregulated and 320 genes were downregulated (Figure 1C).

### 2-AA modulates multiple signaling pathways in HPMEC and NHBE cells

We conducted a pathway enrichment analysis to identify the pathways modulated by 2-AA. The pathways that represented Reactome were organized according to the statistical significance, assessed through p-value and false discovery rate (FDR). Analyses of upregulated and downregulated genes and associated pathways in HPMEC cells are shown in Table 1. Specifically, analysis of the 2558 upregulated genes in these cells revealed 30 enriched pathways primarily linked to proinflammatory response, including interleukins (IL-17, IL-26, IL-3, IL-7), MAPK1/MAPK3 signaling, VEGFA-VEGFR2 signaling and RUNX1 mediated transcriptional regulation (Figure 1D) (43). Concurrently, anti-inflammatory responses, IL-10, IL-4, and IL-13 signaling pathways were also upregulated (Figure 1D). Other upregulated inflammatory responses were associated with the signaling pathways IL-6, TGF-beta, FoxO, and metabolic responses like PI3-Akt, and receptor tyrosine kinase (Figure 1D). In addition, G protein-coupled (GPCR) signaling, known for its contribution to modulating inflammatory cascades and influencing neutrophil degranulation, both critical components of innate immune response (44), was significantly upregulated. Additional upregulated pathways include ESR (estrogen receptor) signaling, nuclear transcription pathway, Hypoxia-inducible factor 1 (HIF-1) signaling, and vitamin D receptor pathway (Figure 1D). Importantly, our analysis showed the upregulation of histone genes and pathways related to the cellular response to stress and degradation of the extracellular matrix, including the expression of matrix metalloproteinases (MMPs) such as *MMP15*, *MMP19*, *MMP25, MMP3*, keratinization, and glycosaminoglycan metabolism (Table S1).

Moreover, the analysis of 1941 downregulated genes in HPMEC cells identified 16 pathways predominantly linked to interferon-alpha/beta/gamma signaling and IL-1 signaling, contributing to a decrease in inflammatory responses (). Along these lines, Toll-like receptor signaling and p53 signaling were downregulated (Figure 1E), a result that might be associated with prolonged (20 hrs) rather than acute exposure to 2-AA. Similarly, 2-AA was found to downregulate the biosynthesis of nucleotide sugars, which are crucial for glycosylation, nicotinate metabolism, and metabolism of water-soluble vitamins and cofactors. In addition, 2-AA leads to a downregulation of oxidative phosphorylation, potentially resulting in mitochondrial dysfunction and increased production of reactive oxygen species (ROS), further influencing inflammation and cell stress response (1, 28). Notably, 2-AA downregulates the metabolism of important homeostatic molecules, including steroids, cholesterol, fatty acids, retinoic acid synthesis, tryptophan metabolism, and phosphatidylinositol signaling (Figure 1E). These findings underscore the significant role of 2-AA in modulating the transcription of multiple genes and pathways that govern endothelial function (Figure 1D).

The Reactome pathways analysis on the effect of 2-AA on bronchial-airway epithelium responses revealed 12 significant pathways and 303 upregulated genes. These pathways were primarily linked to proinflammatory, such as IL-12 and VEGFA-VEGFR2 signaling, and anti-inflammatory responses, such as IL-4 and IL-13 signaling (Figure 1F). Furthermore, pathways related to neutrophil degranulation, cellular response to stress, and keratinization associated with cell survival and differentiation were also upregulated. Notably, and in contrast to the effects on endothelial cells described above, the mitochondrial oxidative phosphorylation pathway exhibited upregulation, as well as other pathways related to the vitamin D receptor pathway and metabolism. The latter included pathways associated explicitly with steroid, cholesterol, lipid, and glycolipid metabolism (Figure 1F). These findings may reflect the different roles of endothelium and epithelium in handling exposure to 2-AA and the progress to the chronic phase of the associated infection.

In contrast, the analysis of the 320 genes that underwent downregulation in NHBE cells revealed significant enrichment in 10 pathways predominantly associated with antibacterial and antiviral responses. These responses were mediated by interferon-alpha/beta/gamma signaling and negative regulators of DDX58/IFIH1 signaling. Additionally, we observed the downregulation of pathways related to toll-like receptor signaling, which is associated with the early innate immune response and TNF signaling (Figure 1G). Also downregulated were pathways related to signaling by receptor tyrosine kinases involved in cell differentiation and the degradation of the extracellular matrix-associated genes crucial for remodelling (45). Notably, the downregulation was extended to pathways involved in the metabolism of nucleotides, steroids, and retinol (Figure 1G). The analysis of both upregulated and downregulated genes and their associated pathway for NHBE cells is shown in Table 2 and is reflective of the developed robust inflammatory response.

In summary, these results indicate that exposure to the quorum-sensing molecule 2-AA induces extensive transcriptional changes in lung endothelial and epithelial cells. The distinct response observed in the two different cell types implies a complex interaction between the bacterial signaling molecule and the specific cellular mechanisms in these cells.

### Insights into the common and different pathways and genes regulated by 2-AA in HPMEC and NHBE cells

The comprehensive pathway analysis of the 2-AA-treated HPMEC and NHBE cells prompted us to determine the common genes and their associated pathways that are significantly regulated by HPMEC and NHBE cells in response to 2-AA. On comparing the 4499 genes from HPMEC and 623 genes from NHBE cells (Table 2), we found a total of 271 genes that were common and significantly regulated in HPMEC and NHBE cells (Figure 2A). A pathway enrichment analysis to determine the pathways and genes that are either upregulated or downregulated found the following genes to be upregulated by 2-AA: *PIM1*, *TIMP1*, *ANXA1*, *HMOX1*, *S1PR1*, which are interconnected with the IL-4 and IL-13 signaling pathway, promoting an anti-inflammatory response. Additionally, genes such as *SH3BGRL3, ANXA1, CCN1, MMRN2, NFATC2, PLAUR, and SIPR1* associated with the proinflammatory response and increased endothelial cell permeability via VEGFA-VEGFR2 signaling were also upregulated. Furthermore, genes *H2BC11, H2BC4, H2BC8, CTSV, and NFATC2* regulated by RUNX1 were upregulated, contributing to cell differentiation and innate immunity. Genes related to keratinization, including *KRT13, KRT16, KRT6C, LCE3D, PI3, PKP1, SPRR1A, SPRR1B, SPRR2A, SPRR2D, SPRR2E, SPRR2G, SPRR3* were similarly upregulated. Other notable upregulated genes included *B3GNT5, FTH1, HMOX1, NAV3, PPARD, SERPINB2,* and *TGFBR3* which are related to the nuclear receptors meta pathway as well as *PRDM1, BMP6, KRT13, KRT16, NFATC2, PPARD, SERPINB1, SPRR1B* associated with the Vitamin D receptor pathway (Figure 2B).

**Figure 2:**
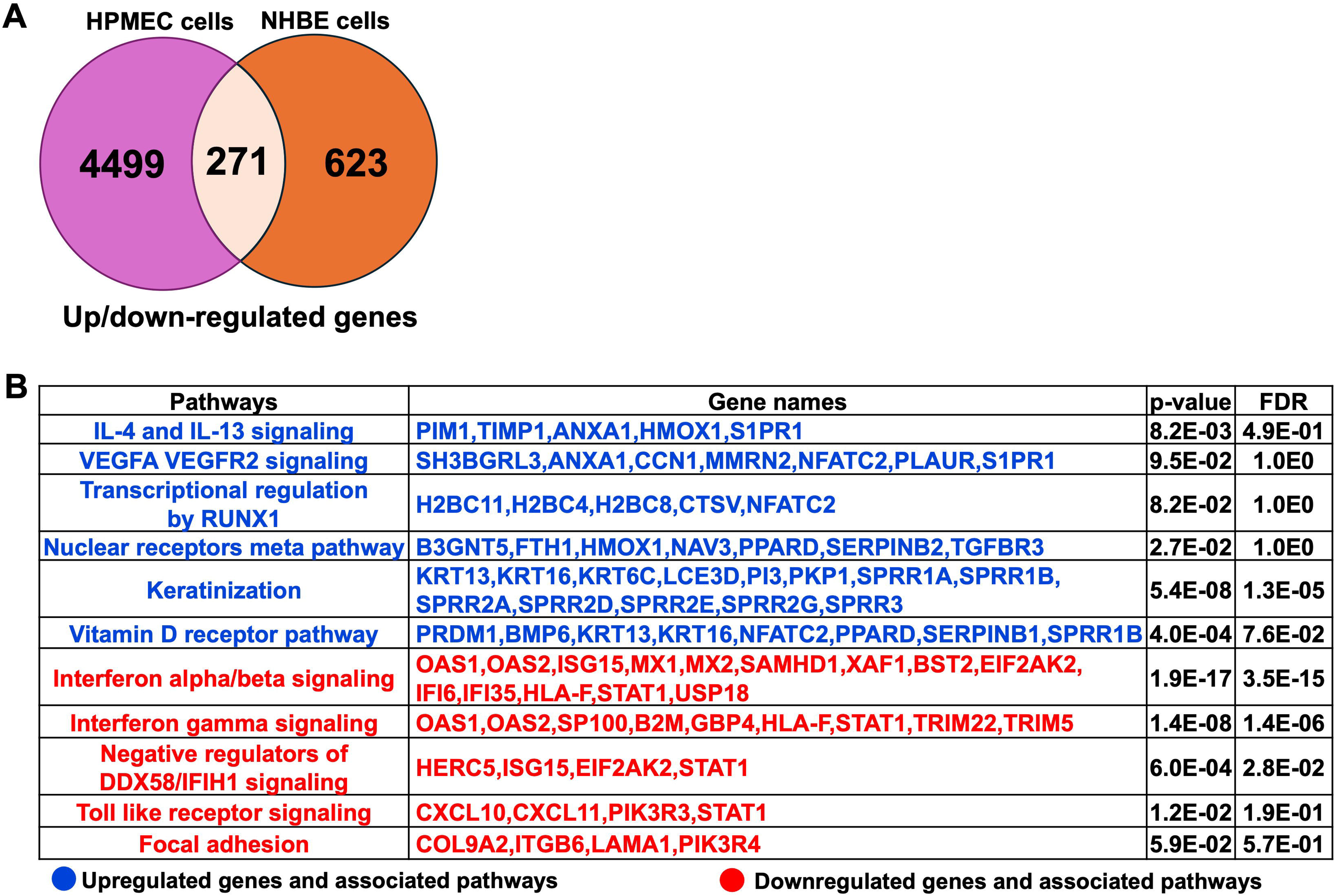
Common genes and associated pathways regulated by 2-AA in HPMEC and NHBE cells. **(A)** Venn diagram showing 271 overlapping and differentially expressed genes between NHBE cells and HPMEC cells identified by RNA seq. **(B)** Upregulated and downregulated genes and their associated pathways are common in NHBE and HPMEC cells. Blue represents upregulated genes and their corresponding pathways, while red represents downregulated genes and corresponding pathways with p<0.05 and a False discovery rate (FDR<0.05). The Reactome pathway enrichment analysis was done using the DAVID web server.

Conversely, the following pathways and genes were downregulated in both cell types: those involved in interferon-alpha/beta signaling (*OAS1, OAS2, ISG15, MX1, MX2, SAMHD1, XAF1, BST2, EIF2AK2, IFI6, IFI35, HLA-F, STAT1, USP18*) and negative regulators of DDX58/IFIH1 signaling (*HERC5, ISG15, EIF2AK2, STAT1*) which are associated with antiviral responses. Additionally, genes related to interferon-gamma signaling (*OAS1, OAS2, SP100, B2M, GBP4, HLA-F, STAT1, TRIM22, TRIM5*) associated with antibacterial response were also downregulated in both the cell types. The Toll-like receptor signaling pathway and downstream genes *CXCL10, CXCL11, PIK3R3, and STAT1* related to innate immune response were similarly downregulated. Furthermore, genes such *as COL9A2, ITGB6, LAMA1,* and *PIK3R4,* which regulate the focal adhesion pathway, were downregulated, indicative of the potential impact on decreased cell migration and proliferation and impaired tissue repair processes(Figure 2B).

Next, genes and associated pathways specifically expressed only in HPMEC or NHBE cells in response to 2-AA are shown in (Table S2). The pathways and genes related to the proinflammatory IL-12 signaling, antiviral response via interferon-alpha/beta signaling, keratinocyte differentiation, steroid metabolism, glycosphingolipid biosynthesis, and pyrimidine metabolism were differentially regulated in NHBE cells (Table S2A). In contrast, the pathways specific to HPMEC cells included signaling pathways regulated by IL-17, IL-3, IL-7, IL-1, IL-18, Il-6, TLR4, IL-10, IL-4, IL-13, TNF, TGFB, FOXO, VEGFA-VEGFR2, PI3-Akt, NF-kappa B, MAPK, receptor tyrosine kinase, ESR, p53, HIF-1 and neutrophil degranulation. Pathways related to lung fibrosis, cellular response to stress, degradation of extracellular matrix, and glycosaminoglycan biosynthesis were also affected by 2-AA in HPMEC cells. Additionally, pathways associated with retinol metabolism, fatty acid metabolism, and phosphatidylinositol signaling were also influenced by the 2-AA treatment in HPMEC cells (Table S2B). Thus, the 2-AA response on HPMEC and NHBE cells underscores the importance of cellular context and experimental setting in determining the potential outcome of inflammatory responses and metabolic processes *in vivo*.

### 2-AA modulates the expression of genes associated with mitochondrial function and cholesterol metabolism in human lung cells

Several of the 271 genes identified as common were differentially expressed in the HPMEC versus NHBE cells. In HPMEC cells, we found upregulation of genes associated with the degradation of extracellular matrix (*COL4A6, COL8A1, FBN2*) by 2-AA, indicating potential alterations in endothelial barrier integrity. In contrast, in NHBE cells, these genes exhibited downregulation, reflecting differing cellular responses that may influence barrier permeability and subsequent cellular functions in the epithelium (Figure 3A).

**Figure 3:**
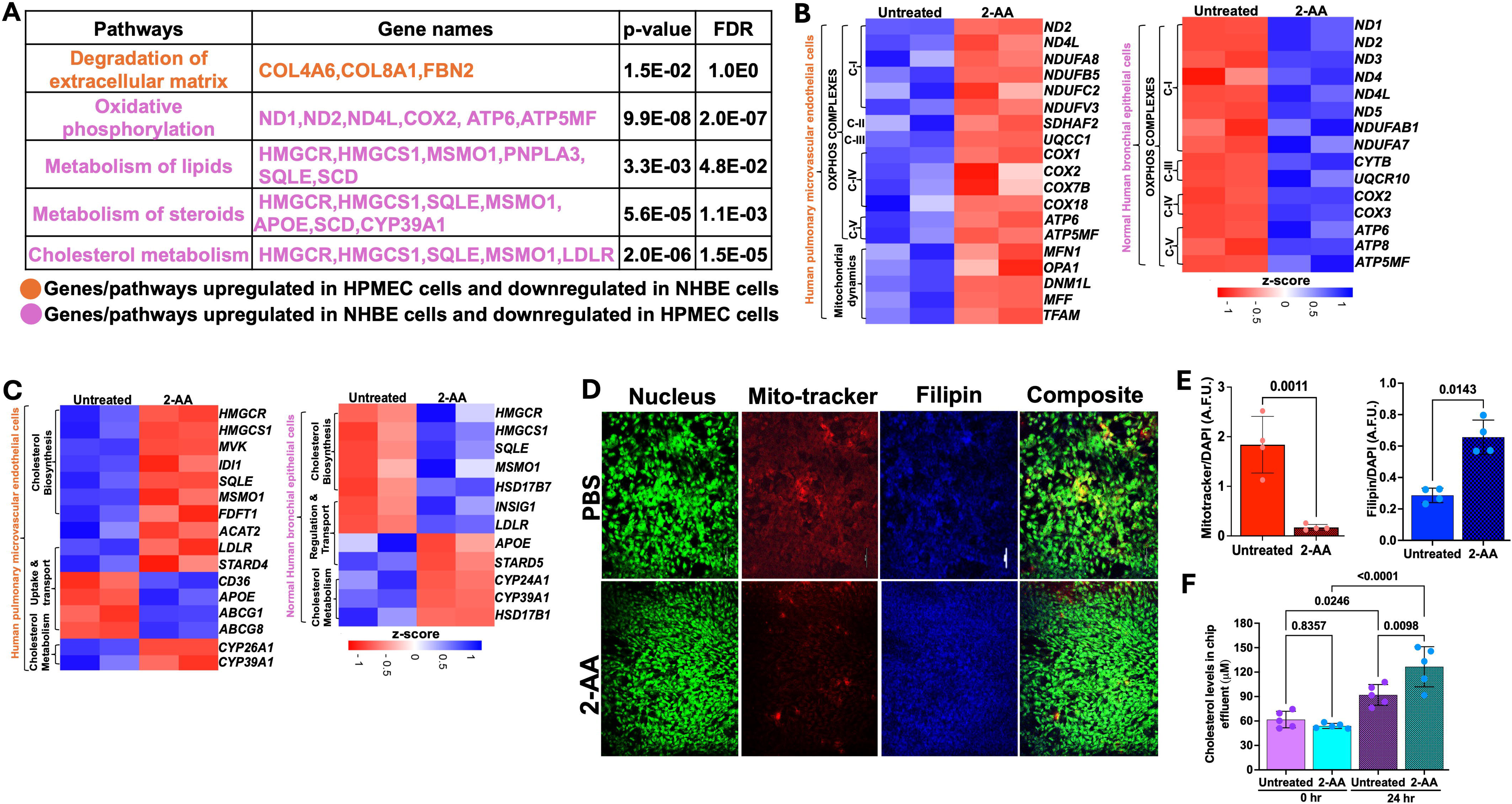
2-AA differentially regulates the mitochondrial and metabolic functions in HPMEC and NHBE cells. **(A)** Table depicting the differentially expressed genes and their associated pathways in the HPMEC and NHBE cells after continuous flow with 20 µM 2-AA for 24 hrs in the lung-on-chip. The orange color represents the genes and pathways upregulated in HPMEC cells and downregulated in NHBE cells. The purple color represents the genes and pathways upregulated in NHBE cells and downregulated in HPMEC cells. Significance is denoted by p<0.05 and FDR<0.05. **(B)** Heatmap showing the differentially expressed genes related to mitochondrial OXPHOS and mitochondrial dynamics in HPMEC and NHBE cells. The Z-score represents the upregulated (blue) and downregulated (red) genes. **(C)** Heatmap showing the differentially expressed genes related to cholesterol biosynthesis, uptake, transport, and metabolism in HPMEC and NHBE cells. The Z-score color scale is consistent with panel B. **(D)** Representative confocal images showing total cholesterol staining with Filipin (blue) and mitochondrial respiration staining with Mitotracker (red) in chips (n=2). Nuclei are counter-stained with DAPI (97). PBS-treated cells were used as a control. The scale bar represents 50 µm. **(E)** Corresponding graphical intensity representing the Arbitrary fluorescence units (A.F.U.) depicting the intensity of mitotracker (red) versus intensity of DAPI (97) and intensity of Filipin (blue) to DAPI (97). (F) Amplex red quantification of total cholesterol from the effluent of the lung-on-chip cells at 0 hrs and 24 hrs either untreated (PBS) or treated with a continuous flow of 20 µM 2-AA. In panel E, each dot represents four independent experiments (n=4); in panel F, each represents five different biological replicates (n=5). The error bars denote mean ± standard deviation. In panel E, p-values were calculated using a two-tailed unpaired t-test, while in panel F, p-values were calculated using ordinary one-way ANOVA followed by Tukey’s post-test. Exact p-values for the pairwise comparisons are depicted on the graphs. The overall p-value summary of panels E and F was p<0.01, indicating statistical significance.

2-AA has been shown to disrupt key mitochondrial functions in skeletal muscle and affect the TCA cycle in macrophages, leading to increased oxidative stress and decreased energy production (Bandopadhaya et al., 2018; Chakraborty et al., 2024). In HPMEC cells, we also found that 2-AA downregulates mitochondrial functions. Genes associated with mitochondrial OXPHOS complex (*ND1*, *ND2, ND4L,COX2, ATP6, ATP5MF*) were downregulated (Figure 3A). The heatmap analysis further emphasizes this effect by showing a consistent decline in the expression of genes encoding mitochondrial complexes I-V alongside essential genes managing mitochondrial dynamics such as *MFN1, OPA1, DNM1L, MFF,* and *TFAM* (Figure 3B).

Conversely, NHBE cells exhibited a distinct response pattern where the genes associated with the mitochondrial OXPHOS complex were upregulated in response to 2-AA (Figure 3B). This suggests an adaptive or compensatory mechanism whereby NHBE cells enhance mitochondrial function in the presence of 2-AA, an additional mechanism of the mounting inflammatory response of the epithelium. The ability of NHBE cells to upregulate mitochondrial complex I and complexes III-V indicates changes in metabolic capability relative to HPMEC cells. This could be linked to differences in cell types, stress responses, or inherent metabolic capacities triggered by inflammation between these cell types. Furthermore, we observed contrasting gene expression patterns relating to lipid, steroid, and cholesterol metabolism in NHBE and HPMEC cells. Notably, NHBE cells exhibited a significant upregulation of several key genes, including *HMGCR, HMGCS1*, *SQLE, MSMO1, APOE, LDLR, SCD, CYP39A1, PNPLA3*, which are integral to lipid biosynthesis and/or storage and cholesterol metabolism (Figure 3A). This indicates lipid synthesis in response to 2-AA in the NHBE cells. In contrast, HPMEC cells downregulated similar cholesterol biosynthesis genes, specifically *HMGCR*, *HMGCS1, MVK, IDI1, SQLE, MSMO1, and FDFT1* (Figure 3C). This reduction suggests a compromised lipid metabolic pathway, impairing cholesterol biosynthesis in the HPMEC cells. In line with the above, we observed a differential regulation of cholesterol uptake and efflux genes. HPMEC cells responded by upregulating *CD36, APOE,* and efflux genes *ABCG1* and *ABCG8,* in contrast to the increase in the expression of the *LDLR* gene found in the NHBE cells (Figure 3C). This suggests cell type-specific alterations in cholesterol biosynthesis, transport, and utilization in support of cell-specific modulation of energy utilization by 2-AA. However, both cell types exhibited downregulation of cytochrome P450 family genes also involved in downstream cholesterol metabolism, specifically *CYP26A1, CYP39A1, CYP24A1, and HSD17B1* that catalyzes the reduction of estrone to the most potent estrogen, estradiol possibly altering receptor signaling (Figure 3C) (46).

### 2-AA induces mitochondrial dysfunction in HPMEC cells and cholesterol accumulation in NHBE cells

Based on the 2-AA mediated differential gene expression of mitochondrial respiration-related genes and cholesterol biosynthesis, uptake, and efflux genes, we stained 2-AA and PBS-treated chips with the Mitotracker dye to label mitochondria and filipin dye to assess for cholesterol abundance in respiring cells (Figure 3D). Fluorescence confocal microscopy images of the stained 2-AA lung-on-chip showed significantly reduced mitochondria staining compared to the PBS-treated control chip, supporting the dysfunction in the bioenergetic capacity suggested in the transcriptomic studies (Figure 3D-E). In contrast, accumulation in cholesterol was observed in the 2-AA-treated cells versus PBS-treated chips (Figure 3D-E). Assessment of the extracellular cholesterol in the effluent collected from the airway-chip HPMEC channel before and after 24 hrs of 2-AA or PBS treatment shows a significant increase in extracellular cholesterol levels at 24 hours compared to the control PBS-treated samples (Figure 3F). These findings indicate a direct correlation between mitochondrial dysfunction and metabolic disturbances.

Cholesterol represents 5-10 % of the lipid component of lung surfactants, playing a vital role in reducing surface tension (47, 48) at the air-liquid interface of the alveoli. The staining results from the lung-on-chip platform cannot differentiate whether the 2-AA mediated accumulation of cholesterol is in the epithelial or endothelial cells. Thus, to determine the cell type associated with the increase in cholesterol observed and correlate the findings with the transcriptome data from the chip suggesting an increase in cholesterol in the epithelial cells, we used chamber slides and stained separately epithelial and endothelial cells, together with evaluation of the cholesterol levels in total cell lysates (Figure 4). First, using the same primary human HPMEC and NHBE cells and fluorescence confocal microscopy, we observed a marked reduction in the Filipin staining in HPMECs treated with 2-AA compared to those treated with PBS (Figure 4A), reflecting a decrease in cellular cholesterol levels in the 2-AA treated cells. Consistent with the microscopy data, quantification of the total cellular cholesterol in 2-AA treated HPMECs cell lysates shows significantly lower amounts of cholesterol relative to PBS-treated controls (Figure 4C). Moreover, the extracellular cholesterol levels from the supernatant of HPMECs before and after 24-hour treatment with 2-AA showed a significant increase in extracellular levels at 24 hours compared to controls (Figure 4E), suggesting alterations in cholesterol synthesis, uptake, or efflux mechanisms due to 2-AA exposure. In contrast, fluorescence confocal microscopy (Figure 4B) of the 2-AA treated NHBE cells stained with Filipin showed an increased intensity of Filipin relative to the PBS controls. To further confirm this finding in other epithelial cells, we measured the total cholesterol content in the human lung epithelial cell line A549, a cell line broadly used in preclinical research (Figure 4D). After 24 hours of 2-AA exposure, A549 cells also demonstrated significantly elevated cholesterol levels compared to PBS-treated cells (Figure 4D). Extracellular cholesterol measurement from these cells’ supernatant collected before and after 24 hours of treatment with 2-AA revealed no significant changes in the extracellular cholesterol levels either between the two time points or when compared to the PBS-treated (control.) These results support the notion that while HPMEC cells actively synthesize and efflux cholesterol, epithelial cells synthesize and accumulate it at extracellular levels, they did not result in measurable changes between 0 and 24 hours (Figure 4F). Further support for these findings is the increase in the lung-on-chip of the *HMGCR* gene expression in epithelial cells (Figure 3C) and the protein levels of HMGCR observed in murine lung section of 2-AA-treated mice stained with the anti-HMGCR antibody against the rate-limiting enzyme HMG-COA reductase (Figure S3A). We observed that 2-AA treated lung tissues exhibited thickness of the epithelial cells compared to the untreated lung tissues (Figure S3A). This finding could be associated with cholesterol accumulation in the lung tissues of 2-AA-treated mice. Moreover, since excessive accumulation of cholesterol results in lung inflammation and fibrosis (49), we performed a hematoxylin and eosin staining (H&E) to investigate whether enhanced cholesterol levels in the lungs are associated with significant lung fibrotic remodeling. Figure S3B shows a progressive thickening of bronchiolar and alveolar walls, indicating structural alterations in lung architecture in the 2-AA treated versus untreated lung tissues.

**Figure 4.**
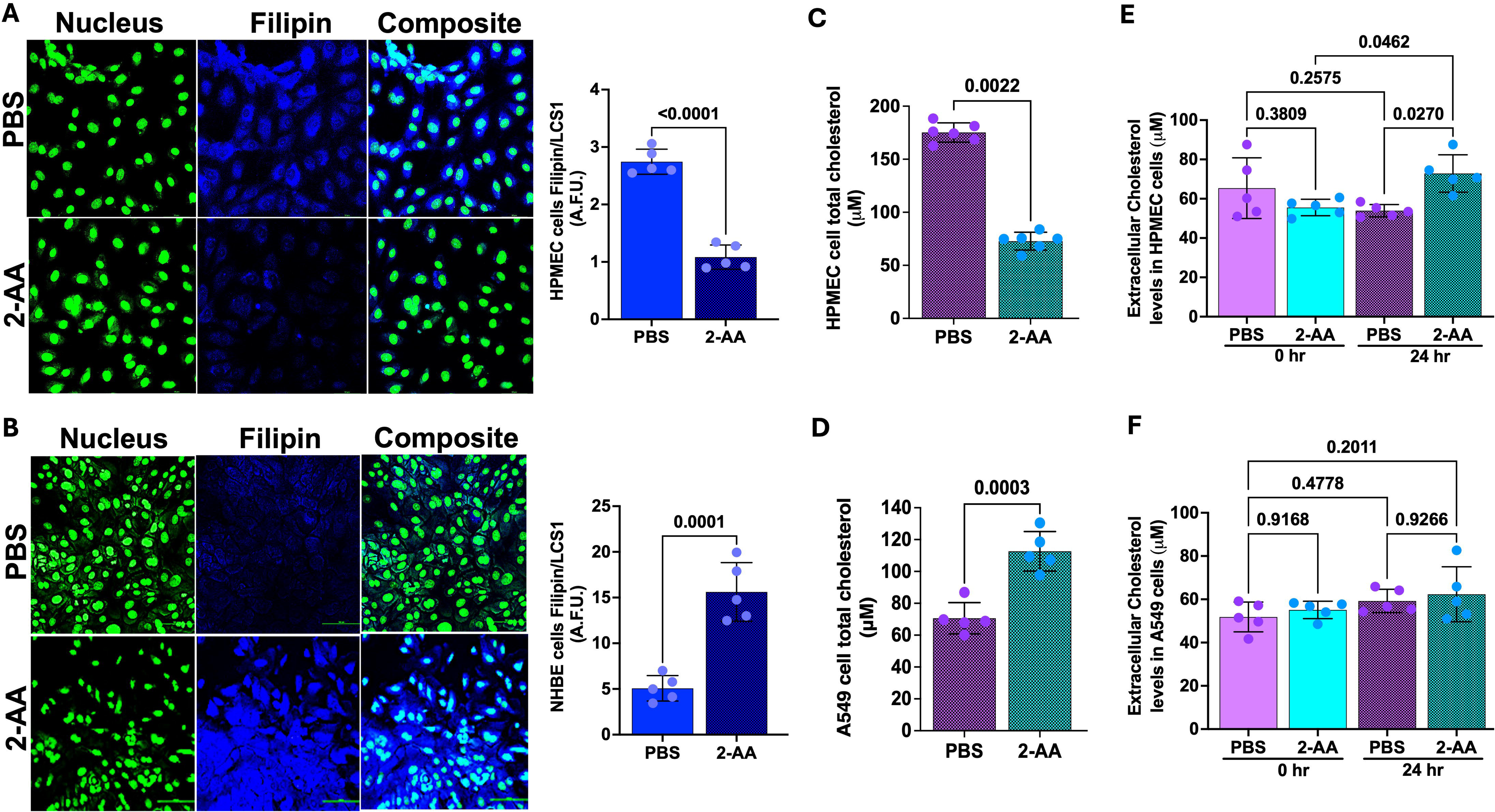
2-AA promotes cholesterol accumulation in epithelial cells. (A-B) Representative confocal images of HPMEC and NHBE cells treated either with 200 µM 2-AA or PBS in chamber slides. Total cell cholesterol stained with Filipin (blue), and nuclei stained with LCS1 (97). PBS was used as a control. Scale bars represent 20 µm. Corresponding graphical intensity plots show arbitrary fluorescent units (A.F.U.) of Filipin (blue) versus LCS1 (97). **(C-D)** Amplex red quantification of total cholesterol in whole cell lysates of HPMEC **(C)** or A549 **(D)** cells 24 hrs post-2-AA treatment (200 µM) (n=5) or PBS in cell culture plates. **(E-F)** Amplex red quantification of total cholesterol in the supernatant of HPMEC **(E)** or A549 **(F)** cells at 0 and 24 hrs, treated with 200 µM of 2-AA of PBS (n=5). Each dot represents five different biological replicates (n=5). The error bars denote the mean ± standard deviation from the average of five samples. In panel (A-D), p-values were calculated by a two-tailed unpaired t-test. In panel (E-F), p-values were calculated by ordinary one-way ANOVA followed by Tukey’s post-test. Exact p-values for the pairwise comparisons are depicted on the graphs. The overall p-value summary of the panel (A-D) is p<0.0001, and of the panel (E-F) is p<0.01, indicating significance.

### 2-AA modulates the expression of cystic fibrosis and idiopathic pulmonary fibrosis-related genes in both HPMEC and NHBE cells

Cystic fibrosis (CF) and idiopathic pulmonary fibrosis (IPF) are complex genetic disorders involving chronic inflammatory changes in the epithelial cells, affecting lung functions (50, 51). Our findings indicate that 2-AA significantly modulates the expression of genes associated with CF (Figure 5A-B and Figure S4) and IPF (Figure 5C-D) in both HPMEC and NHBE cells. Heatmap analysis revealed that *in* HPMEC cells, 2-AA led to the downregulation of cystic fibrosis gene marker *CFTR*, essential for bicarbonate secretion while upregulating *KLF4*. This transcription factor can directly suppress the expression of the *CFTR* gene (52). HPMEC and NHBE cells showed upregulation of the mucin genes (Figure 5A-B), *MUC5B,* and *MUC4,* contributing to mucus accumulation in the lungs. These genes were also associated with IPF (Figure 5C). Additionally, *KCNE3*, a voltage-gated potassium channel critical for normal chloride ion transport (53), was downregulated in response to 2-AA in endothelial cells. At the same time, the H+/K+-ATPase *ATP12A* (54), associated with airway surface liquid (55) and acidification, was upregulated in both cell types.

**Figure 5:**
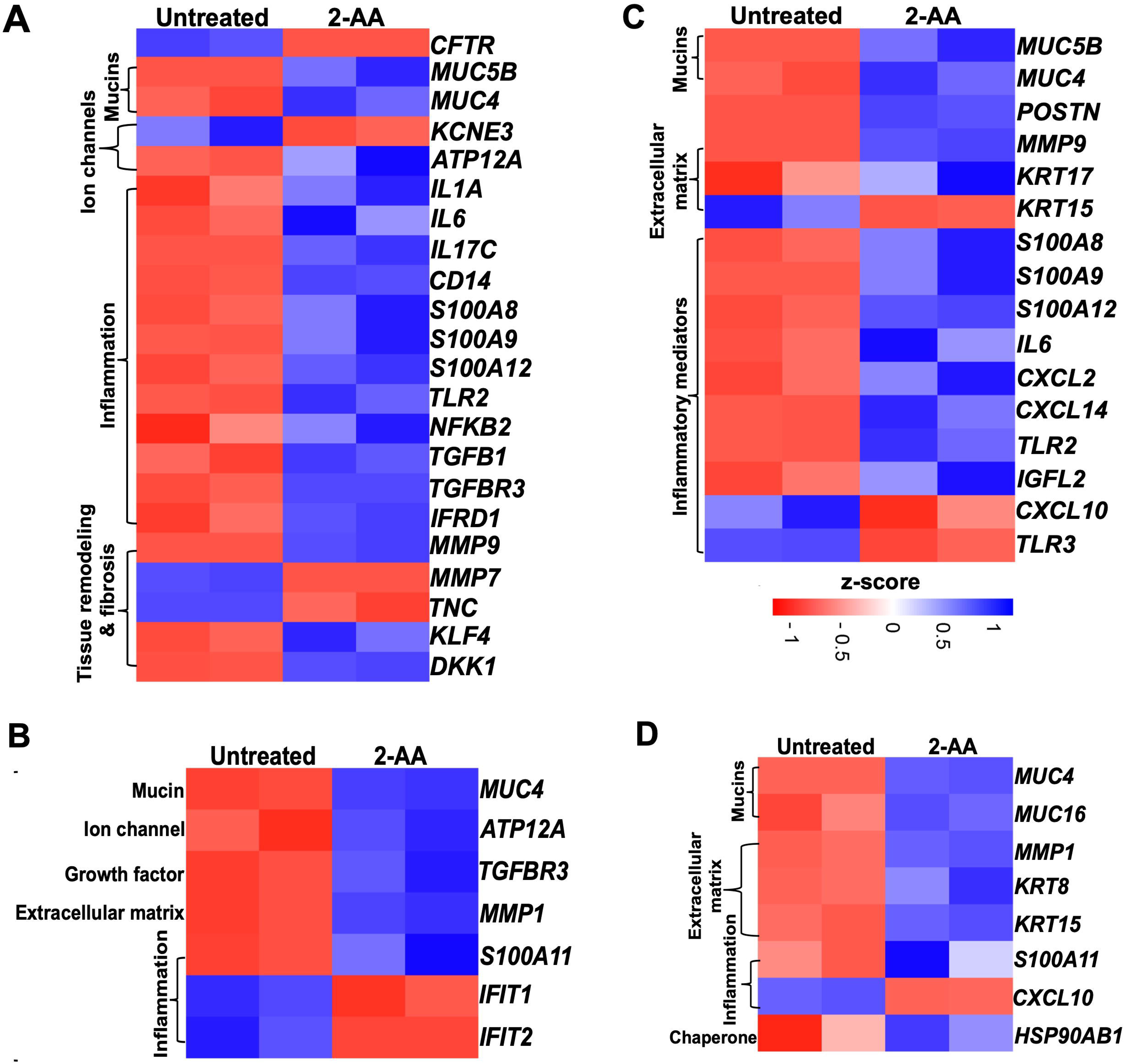
Airway-on-chip studies reveal the impact of 2-AA on the expression of cystic fibrosis and idiopathic pulmonary fibrosis-related genes in HPMEC and NHBE healthy cells. (A-B) Heatmap depicting the gene markers associated with cystic fibrosis in healthy HPMEC and NHBE cells after treatment with 20 µM 2-AA for 20 hrs under continuous flow conditions. **(C-D)** Heatmap showing the expression of gene markers associated with idiopathic pulmonary fibrosis in healthy HPMEC and NHBE cells under the same treatment conditions as (A). z-score color range represents the upregulated (blue) and downregulated (red) genes.

The treatment with 2-AA also induced an unbalanced cytokine response in endothelial cells, with upregulation of proinflammatory cytokines (56–58) such as *IL1A*, *IL6*, *IL17C,* and *NFKB2* which are known to amplify inflammation and innate immune response (59, 60), including *CD14*, *TLR2* (Figure 5A-B). Of these, *IL6 and TLR2* associated with IPF were also found to be upregulated with 2-AA treatment (Figure 5C). Furthermore, the *S100* calcium-binding family gene markers for both CF and IPF, acting as chemokines for neutrophil recruitment to sites of inflammation (61–63), were upregulated with *S100A8*, *S100A9*, *S100A12* in endothelial and S100A11 in epithelial. Notably, the Wnt and TGFB signaling pathways have been identified as crucial players in maintaining Iung epithelial polarization and differentiation in the airways, while TGFB is a main driver of fibrosis too. (64). These proteins also play an important role in the maintenance and homeostasis of tissue matrix. Thus, upregulation of *TGFB1* and *TGFBR3* alongside 2-AA treatment suggests increased fibrotic burdens (65), while *DKK1*, a known Wnt antagonist, indicated potential dysregulation of the Wnt signaling pathway (64). Additionally, *IFRD1* (Interferon-related developmental regulator 1), identified as a modifier of cystic fibrosis lung disease severity, was also upregulated in endothelial cells (66). MMPs (matrix metalloprotease), which are implicated in extracellular matrix turnover, tissue degradation, and repair (), exhibited upregulation of *MMP-9*, while *MMP-7* (67, 68) was downregulated in endothelial and MMP1 in epithelial (69). These changes delineate the membrane architecture and tissue homeostasis to be dysregulated.

In the adjacent NHBE cells (Figure 5B), besides the CF gene markers (*MUC4*, the H+/K+ transporter *ATP12A*, and the inflammatory *TGFBR3)* to mirror the expression observed in HPMEC cells, additional genes were found to be expressed explicitly in these epithelial cells. The matrix metalloprotease *MMP1*, which plays a role in degrading extracellular matrix components (70), was upregulated alongside *S100A11*, another calcium-binding gene (71). Conversely, *IFIT1* and *IFIT2* (72), which are associated with the interferon response, were downregulated, potentially increasing the susceptibility to bacteria. The observed dysregulation of inflammation and mucus production by 2-AA suggests a potential role of this signaling molecule in exacerbating lung fibrosis in CF.

Some additional IPF marker genes upregulated in 2-AA treated HPMEC cells, including *POSTN* (periostin) and *KRT17,* both contributing to extracellular matrix production. Conversely, *KRT15* expression was downregulated, which may contribute to the compromised clearance of damaged cells and consequent triggering of fibrosis (73) (Figure 5C). Furthermore, proinflammatory chemokines *CXCL2* and *CXCL14* (74, 75) were upregulated, while *TLR3* and *CXCL10*, associated with anti-fibrotic properties, were downregulated. Also, expression of *IGFL2* (insulin-like growth factor-like family member 2) secreted from the extracellular matrix was also increased (76). Analysis of the IPF gene markers in the adjacent NHBE cells revealed a response to 2-AA treatment comparable to that seen in HPMEC cells, with upregulation of keratin *KRT15* and downregulation of chemokine *CXCL10* (Figure 5D). Genes were found to be upregulated only in NHBE cells. However, part of the functions found to be affected in HPMEC cells included keratin *KRT8* (77), *MUC16* implicated in the fibrotic processes (78) through its interaction with the TGFB1 and the canonical pathways and the cellular senescence-associated gene *HSP90AB1* (79). These results suggest the 2-AA’s potential role in promoting pathological changes associated with lung fibrosis in the endothelial and epithelial cells.

## Discussion

In this study, using cutting-edge technology that mimics the organotypic microenvironment of the human airway, we uncovered multiple common and cell-specific 2-AA-mediated responses, underscoring the multifaceted role of this *PA* QS secreted small molecule on pulmonary function and physiology. The platform’s emulation of the human hemodynamic environment and physiological biomechanics as established by flow and tissue-relevant stretching allowed the evaluation of the impact of this important MvfR-regulated QS molecule on a dynamic and physiological human-relevant environment built by pulmonary microvascular endothelium and bronchial epithelium cells.

The 2-AA multifaceted effects are in line with this study’s transcriptomic findings with the HPMEC and its adjacent NHBE cells showing multiple common and cell-specific 2-AA-mediated responses affecting various signaling pathways and genes. While the endothelial and bronchial epithelial cells are not in direct contact in the lungs, their crosstalk through paracrine signals, inflammation, and shared microenvironment changes play a critical role in regulating lung function during health and disease (80, 81). The microvascular endothelium may secrete a variety of soluble mediators that can diffuse to the epithelium and affect their function (82), promoting intercellular signaling between these two cell types. The setup of the chip with the two cell types settled in separate channels, enables cell-cell communication and permits the assessment of the overall impact of 2-AA on the gene responses of endothelial and epithelial cells and their potential crosstalk in response to this challenge.

Analyses of the transcriptomic findings enabled us to delineate common and distinct physiological changes in epithelial and endothelial cells. We found that 2-AA in HPMEC cells affects key innate immune response pathways involving MAPKs and the concomitant regulation of proinflammatory and anti-inflammatory cytokines (15). As anticipated, specificproinflammatory responses mediated by IL-17, IL-26, IL-3, IL-7 MAPK1/MAPK3 signaling were upregulated, but major anti-inflammatory responses mediated by IL-10, IL-4, IL-13, PI3-Akt signaling were also upregulated in the 2-AA treated HPMEC cells, potentially as a counterbalancing mechanism to drive the clearance of *PA* lung infection at this early stage of the disease.

Furthermore, the genes and pathways associated with extracellular matrix degradation *COL4A6, COL8A1, and FBN2* were upregulated in NHBE. In contrast, the metalloproteinases *MMP1, MMP15, MMP19, MMP3, MMP9, and MMP25* were upregulated in HPMEC cells, presumably leading to the loss of epithelial-endothelial barrier and damage. This observation aligns with the previously established role of these genes in maintaining the basement membrane structure, which is vital for the maintenance of endothelial cell barrier tightness (83–85). More noteworthy is that 2-AA treated HPMEC cells have compromised mitochondrial respiratory chain, ATP production, and consequently energy metabolism within the cells. This is in accordance with the evidence of mitochondrial dysfunction we reported with this MvfR-regulated molecule (1, 2, 29) and in the pathogenesis and development of lung diseases (86). The downregulation of mitochondrial energy metabolism by 2-AA should lead to functional impairments that adversely affect cellular energy homeostasis in HPMEC cells. Interestingly, while the associated pathways known to respond to hypoxic conditions, such as HIF-1 signaling, were upregulated in HPMEC cells, an opposite effect was found on the bronchial epithelial cells, suggesting functional or even hyperactive mitochondria. These results point to the notion that bacterial infection leads to increased aerobic glycolysis in infected bronchial epithelial cells (87) which fights to survive the infection and maintain airway functionality.

Cholesterol is the major neutral lipid in pulmonary surfactant (88). It is known that alterations in cholesterol metabolism and its implications have a fundamental role in pulmonary surfactant composition, contributing to lung function (89). In a lung-on-chip model of murine alveoli, alveolar surfactant shown to play an essential role in mycobacterial growth (90). Our results show significant alterations in cholesterol metabolism in both HPMEC and NHBE cells, a molecule with multifaceted functionalities in cellular processes. In HPMEC cells, we observed an upregulation of genes associated with cholesterol uptake, including *CD36*, *APOE,* and the efflux genes *ABCG1* and *ABCG8,* suggesting an enhanced requirement or capability for cholesterol utilization/storage in these cells. Conversely, the NHBE cells displayed an increase in expression of the *LDLR* gene, which plays a critical role in cholesterol uptake. This was further confirmed by the observed increased extracellular cholesterol in the supernatant of HPMEC cells and the increased intracellular cholesterol in NHBE cells with the 2-AA treatment. The observed pattern implies a potential uptake of extracellular cholesterol by NHBE cells, possibly stemming from efflux mechanisms activated in HPMEC cells. These results suggest a potential crosstalk in cholesterol metabolism between these cells.

Altered cholesterol homeostasis has been reported to be involved in lung inflammation and injury (91), and its accumulation may interfere with the capacity of the lung surfactant, potentially exacerbating conditions such as pulmonary fibrosis. Our results from HMGCR staining of the lung tissue from mice treated with 2-AA show an accumulation of cholesterol in the epithelial cells and thickness in the epithelial cell membrane, affecting functionality and prompting fibrosis. These findings support our results, indicating that 2-AA mediates cholesterol accumulation, which may contribute to the pathogenesis driven by *PA*.

Results show that 2-AA potentially exacerbates lung fibrosis by modulating CF and IPF key genes involved in mucus production, inflammation, and extracellular matrix remodelling in both endothelial and epithelial cells. Interestingly, in HPMEC cells, the expression of the CF gene marker *CFTR* was lower. In contrast,other CF and IPF markers, *MUC5B, MUC4, and MUC16* genes, were higher, indicating hyper-concentrated mucus, ineffective mucociliary clearance, and impairment of the endothelial membrane barrier (92, 93). Furthermore, 2-AA treatment influenced proinflammatory responses in opposing ways between HPMEC and NHBE cells, suggesting a cell-specific mechanism of action. Studies have reported the critical role of endothelial cells in CF-related pathology, particularly in mediating inflammation and vascular remodelling (92, 94–96). These findings suggest that 2-AA can be a potential biomarker or therapeutic target in chronic lung diseases.

Our findings reveal the previously unreported impact of the MvfR-regulated secreted molecule, 2-AA, on primary female bronchial-airway epithelium and pulmonary endothelium cells. The resulting cellular changes may provide new insights into our understanding of lung *PA* infections in healthy individuals and those with CF and IPF. At the core of the findings in HPMEC and NHBE cells are the observed mitochondrial dysfunction, metabolic alterations, and distinct signaling pathways, providing crucial insights for developing cell-specific targeted countermeasures against *PA*.

## Data availability

All bulk RNA sequencing data have been uploaded on the GEO Expression Omnibus (GEO) database and made publicly available with the accession code GSE290885.

## Contributions

L.G.R. contributed to the conception of the study. L.G.R., S.A., and A.C. contributed to the design of the study. S.A., A.C., and V.K.S. performed experiments. S.A. analysed the RNA sequencing data. L.G.R, S.A., A.C., K.K., and S.L. contributed to data interpretation. All authors have approved the submitted version of the study and their contributions. S.A. and L.G.R. wrote the paper.

## Acknowledgments

The authors thank Roberto Plebani and Kelsey Wheeler for their help in setting up the platform in the Rahme Lab and for initiating part of the experiments, Alexandra Dimitriou for some technical assistance, and Rashmi Richa for her help with the mice lung sectioning. This study was supported by the Massachusetts General Hospital Research Scholar Award and NIH awards R01AI177555 and R01AI134857 to LGR. The funders had no role in study design, data collection, interpretation, or the decision to submit the work for publication.

---

**Figure S1: Principal component (PCA) analysis of untreated and 2-AA treated samples. (A)** PCA plot of human pulmonary microvascular endothelial (HPMEC) cells (n=2). **(B)** PCA plot of normal human bronchial epithelial (NHBE) cells (n=2). A clear separation between untreated (PC1) and 2-AA treated (PC2) is observed. The genes were normalized using DeSeq2.

**Table S1: Reactome pathway enrichment analysis of HPMEC and NHBE cells exposed to 2-AA.** Table showing the upregulated and downregulated pathways with the associated genes in HPMEC and NHBE cells with p<0.05 and a False Discovery Rate (FDR<0.05). The count represents the number of genes associated with a pathway.

**Table S2: Non-common 2-AA regulated genes and associated pathways in HPMEC and NHBE cells.** Table showing Reactome pathway enrichment analysis of upregulated genes (blue) and downregulated genes (red) with their associated pathways after 20 hrs of continuous flow of 20 µM 2-AA treatment in human microvascular endothelial (HPMEC) **(B)** and NHBE **(A)** cells. Significance is denoted by p<0.05 and FDR<0.05. The Reactome pathway enrichment analysis was done using the DAVID web server.

**Figure S3: Representative immunofluorescence microscopy of mice lung tissue sections 4 days post-2-AA treatment shows the accumulation of cholesterol. A.** Cholesterol accumulation was visualized using an antibody against the rate-limiting enzyme HMG-CoA reductase (HMGCR) (red). Nuclei (blue) were counterstained with DAPI. PBS-treated cells were used as a control. The scale bar represents 100 µm. B. Representative images of Haematoxylin and eosin (H&E) staining of mouse lung sections. (A) untreated mice depict normal lung histology. (c) 2-AA-treated mice show thickening of the epithelial cell membrane (c). The black boxes highlight (b) normal bronchial epithelial cells in untreated mice and (d) hypertrophy in the bronchial epithelial cells of 2-AA treated mice. Images represent n=4 mice per group, with 6 tissue sections analyzed per mouse.

**Figure S4: Baseline gene expression of cystic fibrosis biomarkers in NHBE and HPMEC cells exposed to PBS (control).** Table showing in NHBE and HPMEC cells, the differential expression of genes increased (blue) or decreased (red) by at least 1.4-fold change with a p-value<0.05. n=2, * p<0.05, ** p<0.01, *** p<0.001. The cloud-computing Galaxy server (usegalaxy.org) was used for data analysis.

