## Supplementary figures and images for "Revealing the impact of *Pseudomonas aeruginosa* quorum sensing molecule 2’-aminoacetophenone on human bronchial-airway epithelium and pulmonary endothelium using a human airway-on-a-chip"

### Supplementary Figure 1

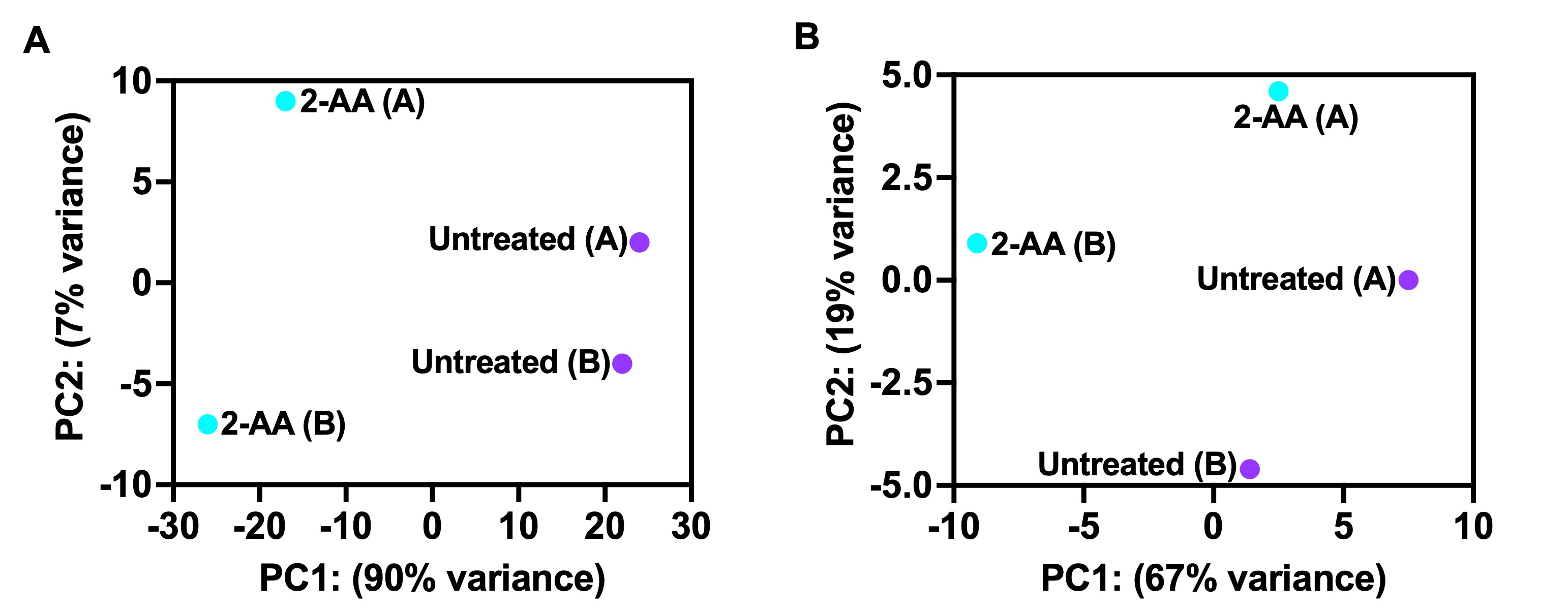

### Supplementary Figure 3

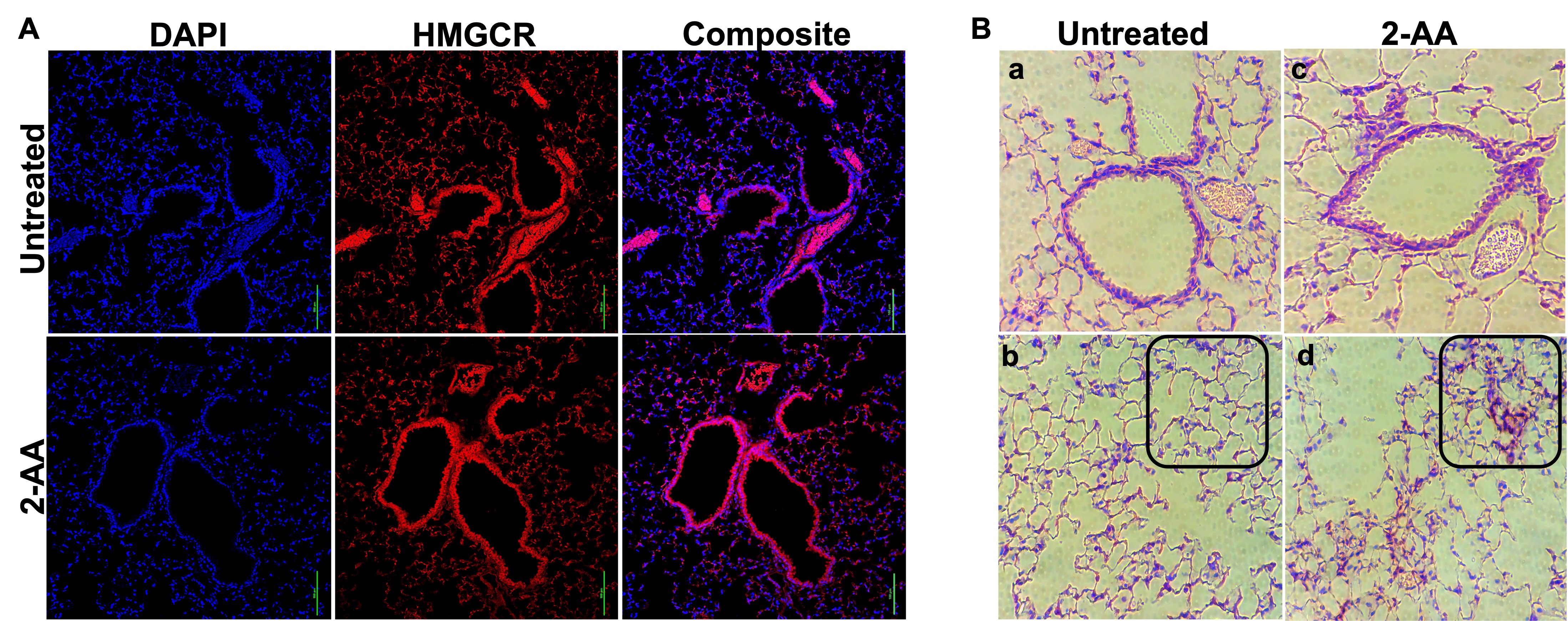

### Supplementary Figure 4

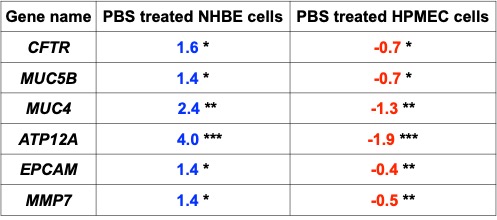
